# Modeling effects of variable preBötzinger Complex network topology and cellular properties on opioid-induced respiratory depression and recovery

**DOI:** 10.1101/2023.08.29.555355

**Authors:** Grant M. Chou, Nicholas E. Bush, Ryan S. Phillips, Nathan A. Baertsch, Kameron Decker Harris

## Abstract

The pre-Bötzinger complex (preBotC), located in the medulla, is the essential rhythm-generating neural network for breathing. The actions of opioids on this network impair its ability to generate robust, rhythmic output, contributing to life-threatening opioid-induced respiratory depression (OIRD). The occurrence of OIRD varies across individuals and internal and external states, increasing the risk of opioid use, yet the mechanisms of this variability are largely unknown. In this study, we utilize a computational model of the preBötC to perform several *in silico* experiments exploring how differences in network topology and the intrinsic properties of preBötC neurons influence the sensitivity of the network rhythm to opioids. We find that rhythms produced by preBötC networks *in silico* exhibit variable responses to simulated opioids, similar to the preBötC network *in vitro*. This variability is primarily due to random differences in network topology and can be manipulated by imposed changes in network connectivity and intrinsic neuronal properties. Our results identify features of the preBötC network that may regulate its susceptibility to opioids.

**Significance Statement:** The neural network in the brainstem that generates the breathing rhythm is disrupted by opioid drugs. However, this response can be surprisingly unpredictable. By constructing computational models of this rhythm-generating network, we illustrate how random differences in the distribution of biophysical properties and connectivity patterns within individual networks can predict their response to opioids, and we show how modulation of these network features can make breathing more susceptible or resistant to the effects of opioids.

## 1 Introduction

Opioid-induced respiratory depression (OIRD) is the primary cause of death associated with opioid overdose. Because both the pain-killing and respiratory depressive effects of opioids require the *μ*-opioid receptor (MOR) encoded by the *Oprm1* gene (Baldo & Rose, 2022; Dahan et al., 2001; Lynch et al., 2023; Sora et al., 1997), there are few effective strategies to protect against OIRD without eliminating the beneficial analgesic effects of opioids. Increasing doses of opioid are often required to maintain analgesia as the neural circuits that mediate pain become desensitized to opioids (Freye & Latasch, 2003; Uniyal et al., 2020), putting patients at greater risk of OIRD. However, a dangerous and less well-understood feature of OIRD is its apparent unpredictability (Dahan et al., 2013). Changes in breathing in response to opioid use can vary substantially between individuals and can be surprisingly inconsistent even within the same individual under different internal and external states or contexts (Cherny et al., 2001; Dahan et al., 2013; Fleming et al., 2015).

Although *Oprm1* is expressed in many brain regions (Erbs et al., 2015), including those involved in the regulation of breathing (Baldo & Rose, 2022; Varga et al., 2020), one site of particular importance is the PreBötzinger Complex (preBötC), a medullary region that is critical for generating the respiratory rhythm (Bachmutsky et al., 2020; Gray et al., 1999; Smith et al., 1991). This network is composed of interacting excitatory and inhibitory interneurons (Baertsch et al., 2018; Harris et al., 2017; Winter et al., 2009). Although inhibitory neurons are important for regulating the frequency and regularity of breathing (Baertsch et al., 2018; Sherman et al., 2015), GABAergic or glycinergic mechanisms do not seem to play a significant role in OIRD in the preBötC (Bachmutsky et al., 2020; Gray et al., 1999). Instead, glutamatergic neurons are the critical component of the preBötC network needed for both rhythmogenesis and OIRD (Bachmutsky et al., 2020; Funk et al., 1993; Greer et al., 1991; Sun et al., 2019). Among the estimated 40-60% of preBötC neurons that express *Oprm1* (Baertsch et al., 2021; Gray et al., 1999; Rousseau et al., 2023), activation of MORs has two primary consequences: Excitability is suppressed due to activation of a hyperpolarizing current, and the strength of excitatory synaptic interactions is reduced (Baertsch et al., 2021). Together, these mechanisms of opioid action act synergistically to undermine the cellular and network mechanisms that mediate preBötC rhythmogenesis.

Neurons in the preBötC have heterogeneous cellular properties, which are readily observed following pharmacological blockade of synaptic interactions. Under these conditions, the intrinsic activity of preBötC neurons is either silent, bursting, or tonic, which largely depends on their persistent sodium conductance (*g*_NaP_) and potassium dominated leak conductance (*g*_leak_) (Butera et al., 1999b; Del Negro et al., 2002; Koizumi & Smith, 2008; Phillips & Rubin, 2019; Yamanishi et al., 2018). However, *g*_NaP_, *g*_leak_, and the intrinsic activity of preBötC neurons are not fixed but can be dynamically modulated by conditional factors such as neuromodulation and changes in excitability (Del Negro et al., 2001; Doi & Ramirez, 2008; Ramirez et al., 2011). Thus, unlike the discrete activity states of its constituent neurons, when synaptically coupled, the network collectively produces an inspiratory rhythm that can operate along a continuum of states as the ratios of silent, bursting, and tonic neurons change (Burgraff et al., 2021; Butera et al., 1999a). As previously demonstrated in rhythmic brainstem slices, the preBötC has an optimal configuration of cellular and network properties that results in a maximally stable inspiratory rhythm. These properties are dynamic, and the state of each individual preBötC network relative to its optimal configuration can predict how susceptible rhythmogenesis is to opioids (Burgraff et al., 2021).

Here, we expand on these findings by utilizing computational modeling to perform preBötC network manipulations and analyses that are experimentally intractable to better understand properties of the network that may contribute to the variation in OIRD and to provide proof of concept for perturbations that may render preBötC rhythmogenesis less vulnerable to opioids. We demonstrate that model preBötC networks exhibit variable responses to simulated opioids. This variation in opioid response is best predicted by differences in “fixed” properties of randomly generated networks, specifically the connectivity between different groups of excitatory and inhibitory neurons as well as which neurons in the network express MOR. In contrast, opioid-induced changes in the intrinsic spiking patterns preBötC neurons (silent, bursting, tonic) do not predict this variation. In networks with high opioid sensitivity, we find that modulation of either *g*_NaP_ or *g*_leak_ can render rhythmogenesis more resistant to opioids. These insights help establish a conceptual framework for understanding how the fixed and dynamic properties of the preBötC shape how this vital network responds when challenged with opioids.

## 2 Methods

### 2.1 Computational modeling of OIRD in the PreBötC

We model the preBötC network as a random, directed graph of *N* = 300 nodes, with each node representing a neuron. The dynamical neuron equations are modified from (Baertsch et al., 2021; Butera et al., 1999a; Butera et al., 1999b; Harris et al., 2017). First, we have the total membrane current balance equation

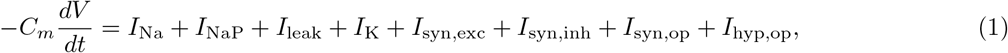

where the currents are given by

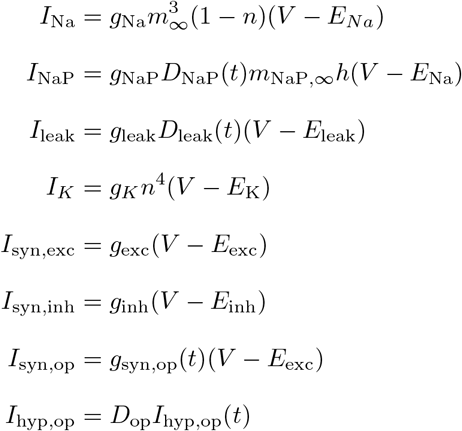

We implemented the terms *D*_NaP_(*t*) and *D*_leak_(*t*) to simulate time-dependent “drugs” strengthening or weakening NaP and leak channel conductances by varying between 0 and 1. We also added the opioid-modulated synaptic and hyperpolarizing currents *I*_syn,op_ and *I*_hyp,op_ as a mechanism for biophysical perturbations through changing *I*_hyp,op_(*t*) and *g*_syn,op_(*t*). While many terms in these differential equations are time-dependent, we only explicitly highlight the time-dependence of *D*_NaP_, *D*_leak_, *g*_syn,op_, *I*_hyp,op_ because these are set exogenously. The gating equations are

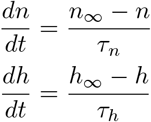

with voltage-dependent steady states,

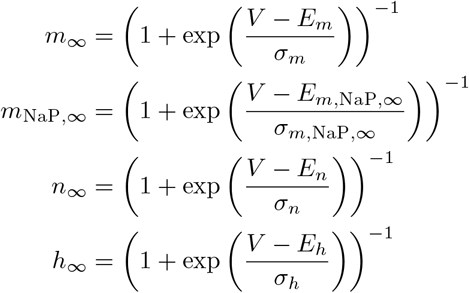

and time constants

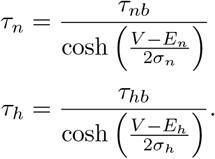

Finally, synapses are modeled separately for excitatory, inhibitory, and opioid-sensitive presynaptic neurons. Each synapse conductance *s* is modeled with first-order dynamics:

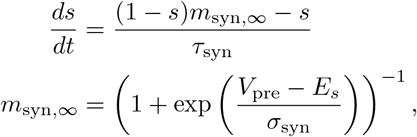

and *g*_syn_ is given as the sum of *g*_syn,max_ · *s* over all incoming synapses to the neuron. This model was implemented in `brian2` (Stimberg et al., 2019). The parameters which are shared across all neurons are given in Table 1; other parameters will be described in the rest of the methods. Our code will be included with the final version of this paper.

**Table 1:**
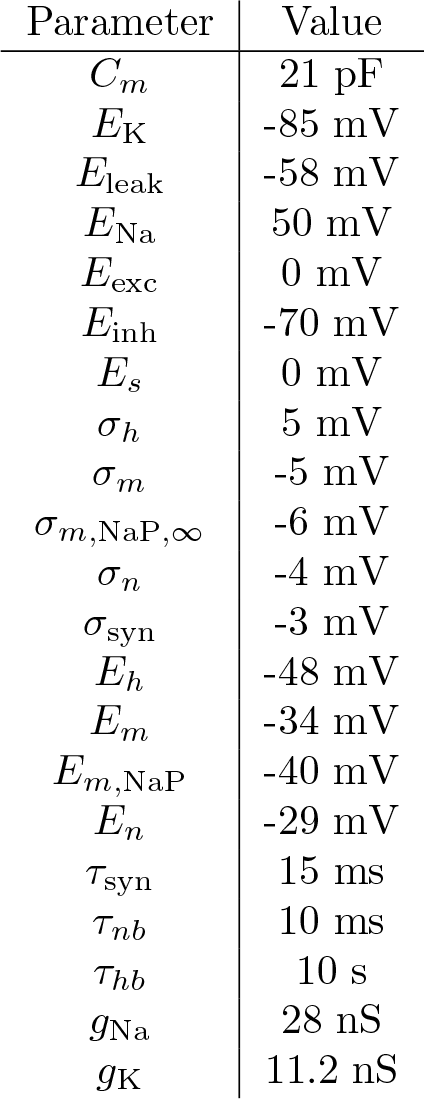
Table of model parameters shared across neurons.

#### 2.1.1 Network construction

Our 300 neuron network consists of 60 inhibitory neurons and 240 excitatory neurons. Synapses were randomly constructed, with each neuron having a connection probability of 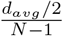, where *d*_*avg*_ is the neurons’ average degree (in-degree + out-degree). Our default *d*_*avg*_ is 6, giving us a connection density of approximately 1%. However, in 3 we increase the connection probability by scaling *d*_*avg*_, e.g. *d*_*avg*_ = 12 results in a 2% connection density.

The intrinsic activity of each neuron is either tonic spiking (T), bursting (B), or quiescent (Q), which is controlled by *g*_leak_ and *g*_NaP_. The *g*_leak_ value for each neuron was randomly drawn from a mixture of three Gaussians with weights [0.35, 0.1, 0.55], means [0.5, 0.7, 1.2] nS, and standard deviation 0.05 nS. The *g*_NaP_ values are drawn from a Gaussian with mean 0.8 nS and standard deviation 0.05 nS. Classification of intrinsic activity is done by using peak detection on the voltage *V* recorded with synapses blocked.

#### 2.1.2 MOR targeting

In all simulations, half of the excitatory neurons are opioid-sensitive (MOR+) and can be targeted with opioid (*D*_op_ = 1), while the inhibitory neurons and the other half of the excitatory neurons (MOR-) are insensitive (*D*_op_ = 0). The opioid-sensitive population’s excitatory synapses follow the *I*_syn,op_ equation, whereas the insenstive neurons follow *I*_syn,exc_. Assignment of *D*_op_ among excitatory neurons is random except in two cases shown in Fig. 5, where opioid is applied to the excitatory neurons with *g*_leak_ values below or above the median among the excitatory population.

#### 2.1.3 Gradual ramp up of opioid

For opioid ramping simulations (Figs. 1, 3, 4E), *I*_hyp,op_ ramped from 0–8 pA, increasing by 0.5% every 3 seconds, while *g*_syn,op_ gradually decreased from 1.0–0.0 nS by 0.5% every 3 seconds. Hence, the total length of the simulation is 10 minutes. The opioid shutdown dosage was calculated by averaging the *I*_hyp,op_ values at the time of the last bursts with amplitudes of 10–15 Hz.

**Figure 1:**
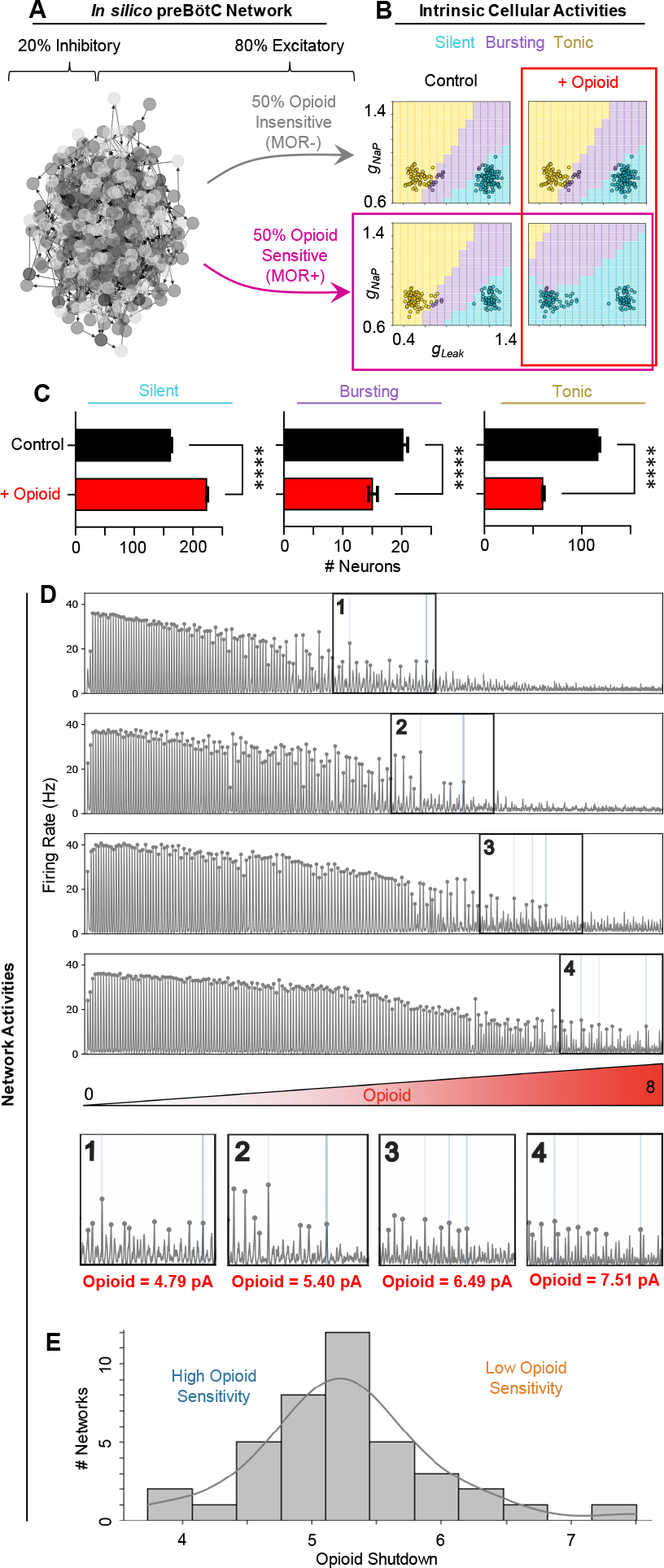
Opioid sensitivity varies across model preBötC networks. (A) A network graph of the *in-silico* preBötC network. (B) Phase diagrams showing intrinsic activities of each neuron (open circles) based on *g*_leak_ and *g*_NaP_ conductances. Top-row: MOR-neurons. Bottom-left: control condition, MOR+ neurons. Bottom-right: opioid applied to MOR+ neurons. (C) Quantified changes in the number of neurons with silent, bursting, or tonic intrinsic activities in response to opioid (*n* = 40 networks; two-tailed paired t-tests; \*\*\*\**p <* 0.0001). (D) Traces of 4 network simulations where opioid is ramped up. Numbered boxes show the last bursts detected at a given amplitude threshold (10-15 Hz/cell). (E) Histogram and kernel density estimation of the distribution of opioid shutdown doses for *n* = 40 model networks.

#### 2.1.4 Timed all-or-nothing perturbations

In simulations with timed all-or-nothing perturbations (Figs. 2, 5A,B,C, 6, and 7), we allowed for a 10 second transient period before each perturbation. Data from transient periods is not used in our analysis. When the opioid perturbation is turned on, *I*_hyp,op_ = 4 pA and *g*_syn,opioid_ = 0.5 nS. We varied *g*_NaP_ (Fig. 6) or *g*_leak_ (Fig. 7). For each node, *g*_NaP_ was increased to 110%, 130%, and 150% of control values, whereas *g*_leak_ was decreased to 90%, 70%, and 50% of control values. In Figs 6 and 7, the 200 s experimental procedure is as follows:

**Figure 2:**
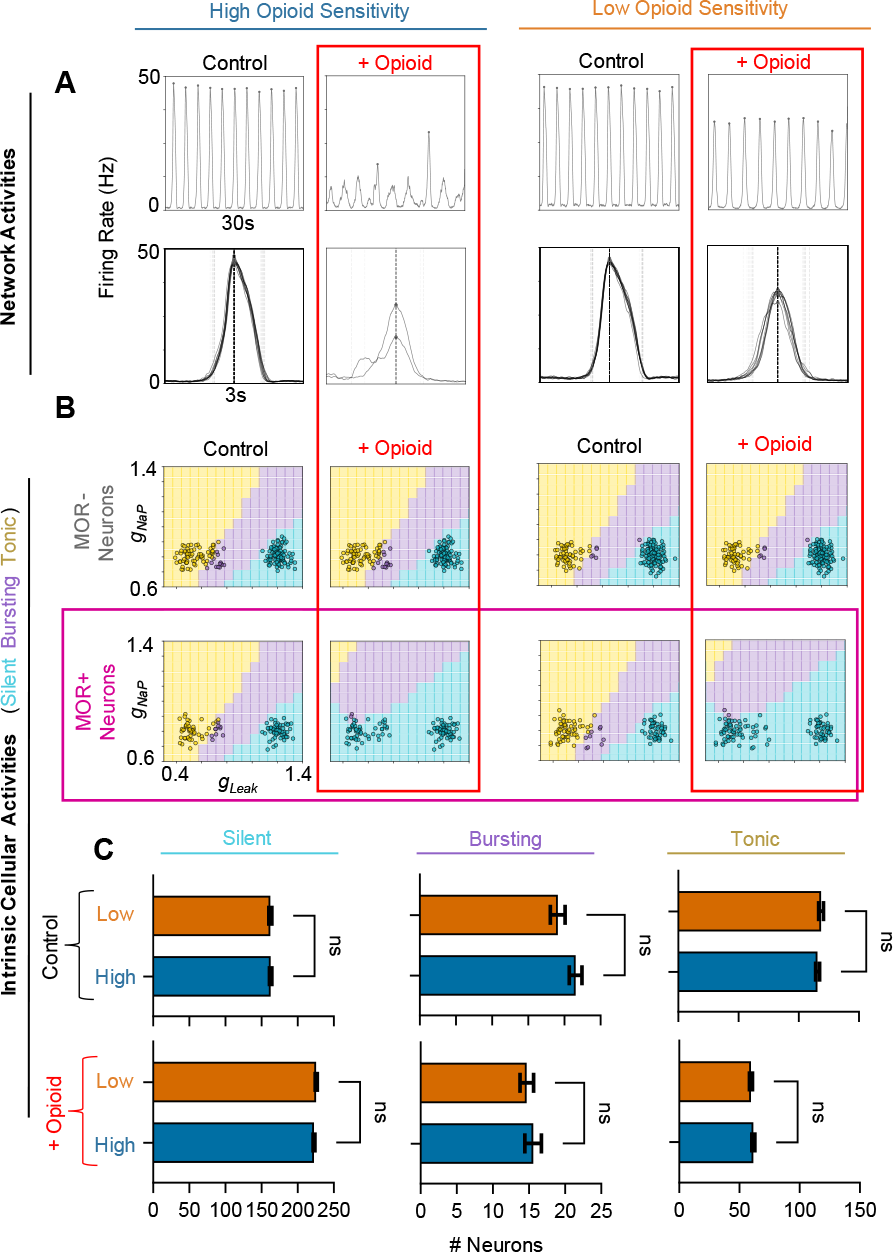
Intrinsic cellular activities do not predict opioid sensitivity. (A) Example rhythms (top) and overlaid burst waveforms (bottom) under control conditions and in the presence of opioid from representative “high sensitivity” (left) and “low sensitivity” (right) networks. (B) Phase diagrams of high (left) and low (right) sensitivity networks, showing intrinsic activities of MOR- (top) and MOR+ (bottom) neurons (open circles) based on *g*_leak_ and *g*_NaP_ conductances. (C) Quantified relationship between opioid shutdown and the number of silent, bursting, and tonic neurons under control conditions (top) and in the presence of opioid (bottom) (*n* = 40 networks; two-tailed paired t-tests; ns=not significant).

1. 10 s transient period
2. 30 s control period
3. 10 s transient period
4. 30 s opioid perturbation
5. 10 s transient period
6. 30 s control “wash” period
7. 10 s transient period
8. 30 s *g*_NaP_ or *g*_leak_ perturbation
9. 10 s transient period
10. 30 s simultaneous perturbation of opioid and *g*_NaP_ or *g*_leak_

### 2.2 Analysis

#### 2.2.1 Burst detection and opioid shutdown dosage

Bursts were detected using a basic peak-finding algorithm (find peaks function in scipy (Virtanen et al., 2020)) where each peak must have a minimum height of 4 Hz/cell and minimum prominence of 10 Hz/cell. We then compute the opioid shutdown dosage by finding the averaging *I*_*opioid*_ values at the time of the last bursts meeting an amplitude threshold of 10, 11, 12, 13, 14, and 15 Hz/cell. Statistical analysis of measured variables was performed using GraphPad Prism 10 software, and data was visualized using a combination of python, GraphPad Prism, and Powerpoint.

#### 2.2.2 Phase diagrams

Each phase boundary was computed by simulating the network under synaptic block, sweeping across a grid of conductances *g*_leak_ ∈ {0.2, 0.3, …, 1.5} nS and *g*_NaP_ = {0.6, 0.7, …, 1.5} nS. The points plotted within the phase boundaries represent the neurons in the two-population network simulated under synaptic block. Population neurons and phase sections are both colored by intrinsic activity classified as described in Section 2.2.1.

### 2.3 Code availability

Code is available in our github repository or upon request to the corresponding author. Code was run using Linux systems with python simulation stack.

## 3 Results

### 3.1 Opioid sensitivity varies across model preBötC networks

In sparsely connected (1%) preBötC networks, connections were drawn randomly between excitatory and inhibitory neurons with different intrinsic activities as determined by varied *g*_NaP_ and *g*_leak_ values. We implemented a two-population distribution of *g*_leak_ and *g*_NaP_ (Baertsch et al., 2021) as described in Methods 2.1.1 to reduce the number of neurons in the network that exhibit intrinsic bursting to 5 − 10% (Fig. 1), as estimated by experimental recordings *in vitro*. With 50% of excitatory neurons randomly designated as opioid-responsive (MOR+) and the remaining 50% non-responsive (MOR-), we performed simulations on 40 different preBötC networks where the effects of opioid (i.e. presynaptic suppression and hyperpolerization) were gradually increased over the course of 10 minutes. An example of how simulated opioid affected the intrinsic activities of MOR+ and MOR-neurons in *g*_leak_, *g*_NaP_ parameter space is shown in Fig. 1B. Across all networks, opioids transitioned the intrinsic activity of preBötC neurons from tonic or bursting to silent (Fig. 1C), similar to observations in preBötC slices (Burgraff et al., 2021). Some networks were highly sensitive to opioid, as shown by a quicker decline in the respiratory rhythm (e.g. Fig. 1D, traces 1 and 2), while other networks were quite resistant (e.g. Fig. 1D, traces 3 and 4). This variability is reflected in the distribution of shutdown dosages, which ranged from 3.73 to 7.51 with a mean of 5.26 and was approximately Gaussian Fig. 1E. Thus, despite all simulated networks having the same number of neurons designated as excitatory, inhibitory, and opioid sensitive (MOR+), there was considerable variation in how individual networks responded to opioids.

### 3.2 Changes in intrinsic cell activities do not predict opioid sensitivity

To explore how random differences in the intrinsic cellular activities of the networks may predict the varied responses of the network rhythm to opioids, we compared networks with high and low opioid sensitivity. “High-sensitivity” networks were defined as those with an above-median opioid shutdown dosage, while networks with a below-median shutdown dosage were considered “low-sensitivity”. Rather than the gradual opioid ramping as shown in Fig. 1, in Fig. 2 we instead simulated a 30-second control period followed by a 30-second period with a moderate dose of opioid applied (opioid=4). The variation in opioid sensitivity is exemplified in Fig. 2A, where we see clear differences in how the rhythm responded to opioid. In the high-sensitivity case, the rhythm became weak and irregular, whereas rhythms produced by the most resistant networks were able to maintain consistent frequencies and burst amplitudes close to baseline. Changes in the intrinsic cellular activities of these representative high- and low-sensitivity networks are shown in *g*_leak_, *g*_NaP_ parameter space in Fig. 2B. Under control conditions and in the presence of opioid, the proportions of neurons with silent, bursting, or tonic intrinsic activity were similar between high- and low-sensitivity networks (Fig. 2C). Indeed, regardless of opioid sensitivity, a similar number of MOR+ neurons that were tonic or bursting in control conditions became silent in the presence of opioid, which was consistent across all 40 networks. Thus, differences in how opioids affect the intrinsic activities of neurons in our model networks are unlikely to explain their variable responses to opioids.

### 3.3 Connection density and network structure regulate opioid sensitivity

To test how the total amount of connectivity with the preBötC model networks affects how they respond to opioids, we ran simulations where the connection probability of each neuron was increased to 2%, 4%, 8%, or 16% for 40 networks each (the default for all other experiments is 1% connection density), while maintaining total synaptic strength in the network constant. The results are shown in Fig. 3. For each trace in Fig. 3A, we can see that networks with higher connectivity are able to maintain a network rhythm at higher doses of opioid. The distribution of dosages that effectively shut down each network also tends to be slightly less variable at higher connection probabilities (3B and C). Thus, preBötC networks with higher total connection densities are more resistant to opioids.

**Figure 3:**
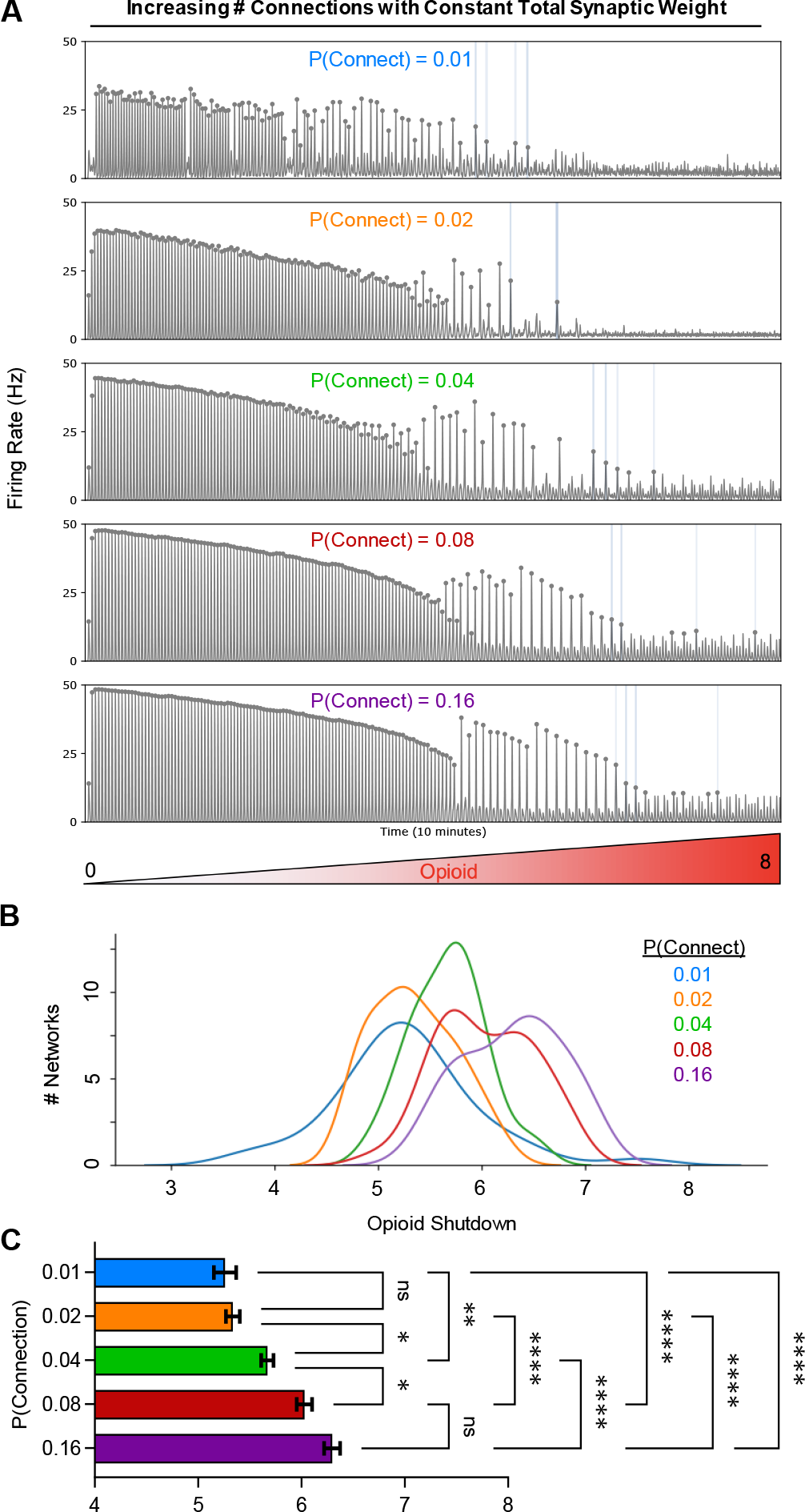
Increased connection density reduces opioid sensitivity. (A) Example traces of 4 different simulations with varied connection densities where opioid is ramped up (opioid=0-8) over 10-minutes. (B) Kernel density estimations showing the distribution of shutdown dosages based on connection probabilities. (C) Quantified opioid shutdown dose vs. connection probability (*n* = 40 networks; one-way RM ANOVA with Bonferroni multiple comparisons tests; *p *<* 0.05, **p *<* 0.01, ****p *<* 0.0001).

Next, we examined how random differences in connection topology may contribute to the variation in opioid responses observed across our 40 randomly drawn model networks. To do so, we first tested whether the total number of excitatory and inhibitory connections (excitation/inhibition balance) within each model network was related to its sensitivity to opioids (Fig. 4A). Correlation analysis revealed that, in general, networks with a more highly connected excitatory population and fewer inhibitory inputs to these excitatory neurons were more resistant to opioids (i.e. higher opioid shutdown dose). In contrast, overall connectivity within the inhibitory population or from excitatory to inhibitory neurons was not correlated with the sensitivity of the network rhythm to opioids. Next, we tested more specifically whether the number of connections within and between, excitatory MOR+, excitatory MOR-, and inhibitory neurons was correlated with the opioid dose that shutdown rhythm generation (Fig. 4B). We found that when the population of excitatory MOR-neurons was more interconnected and received less inhibitory input, the network was more likely to be resistant to opioids.

**Figure 4:**
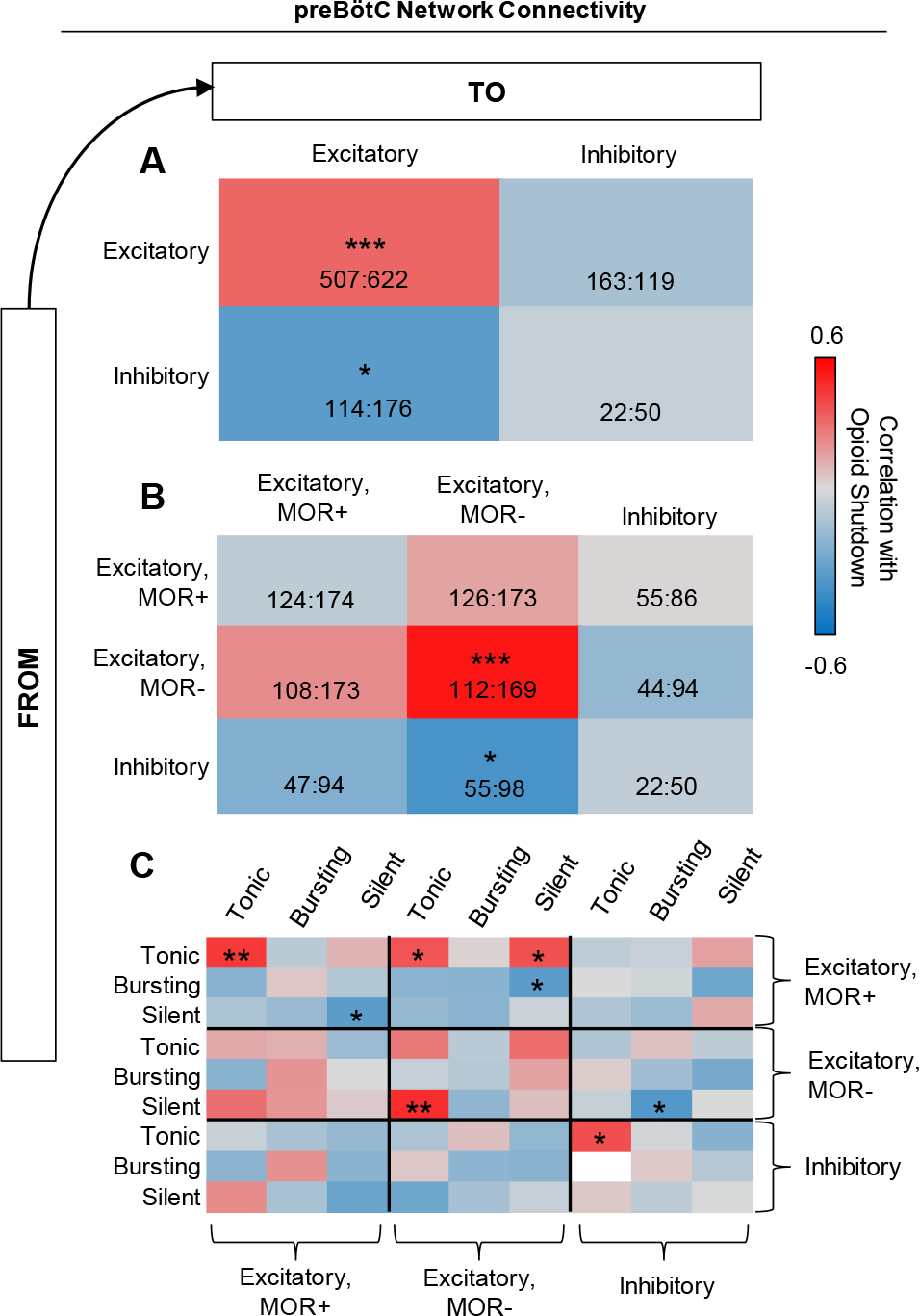
Network structure regulates opioid sensitivity. Correlation analysis of the relationship between opioid shutdown dose and connectivity within and between (A) excitatory and inhibitory populations, (B), MOR+, MOR-, and inhibitory populations, and (C) tonic, bursting, and silent excitatory and inhibitory subpopulations (*n* = 40, two-tailed correlation analysis; *p *<* 0.05, **p *<* 0.01, ***p *<* 0.001). Numbers in A and B represent the max and min number of each type of connection.

In a third analysis, we broke the network connectivity down even further by computing correlations between opioid shutdown dose and the number of connections among intrinsically tonic, bursting, and silent excitatory MOR+, excitatory MOR-, and inhibitory neuron subpopulations (Fig. 4C). This revealed three primary observations. First, the number of connections from silent to tonic excitatory MOR-neurons was the strongest driver of opioid resistance among this MOR-population. Second, although the total number of connections within the MOR+ population was not predictive of opioid sensitivity, networks with more connections between intrinsically tonic MOR+ neurons and fewer connections between intrinsically silent MOR+ neurons were more resistant to opioids. And third, networks were also more likely to be resistant to opioids if they had more connections from tonic MOR+ neurons to tonic or silent MOR-neurons and fewer connections from bursting MOR+ neurons to silent MOR-neurons. Overall, these correlation analyses suggest that differences in network topology as a result of randomness in the assignment of network connections contribute to the variable responses of preBötC networks to opioids.

### 3.4 Identity of MOR+ neurons regulates opioid sensitivity

Because 50% of the excitatory neurons in our model networks are randomly designated as MOR+, we next wondered how the opioid sensitivity of the model networks may be altered if the identity of MOR+ neurons is non-random. To address this question, we performed simulations to compare opioid responses in networks where the intrinsically silent neurons (high *g*_leak_) or the tonic/bursting neurons (low *g*_leak_) were designated as MOR+, as described in Section 2.1.2. Example network activity during these experiments is shown in Fig. 5A. Compared to random assignment of MOR as described above, assigning MOR to the low *g*_leak_ population made the rhythms more resistant to opioids, whereas assigning MOR to the high *g*_leak_ population made them more sensitive. When low *g*_leak_ (primarily intrinsically tonic/bursting) neurons are MOR+, 92.3% of the population became intrinsically silent in response to opioids (Fig. 5B). However, this only reflects the intrinsic activity of those neurons with synapses blocked. With synaptic interactions intact, the network remained rhythmic, with a slower frequency than control conditions but a similar amplitude. On the other hand, when the high *g*_leak_ (primarily intrinsically silent) population is MOR+, the rhythm collapsed under only a moderate dose of opioid (*I*_hyp,op_ = 4 pA) (Fig. 5A). In this case, changes in the intrinsic activities of neurons in the network in response to opioid were minimal (Fig. 5B).

**Figure 5:**
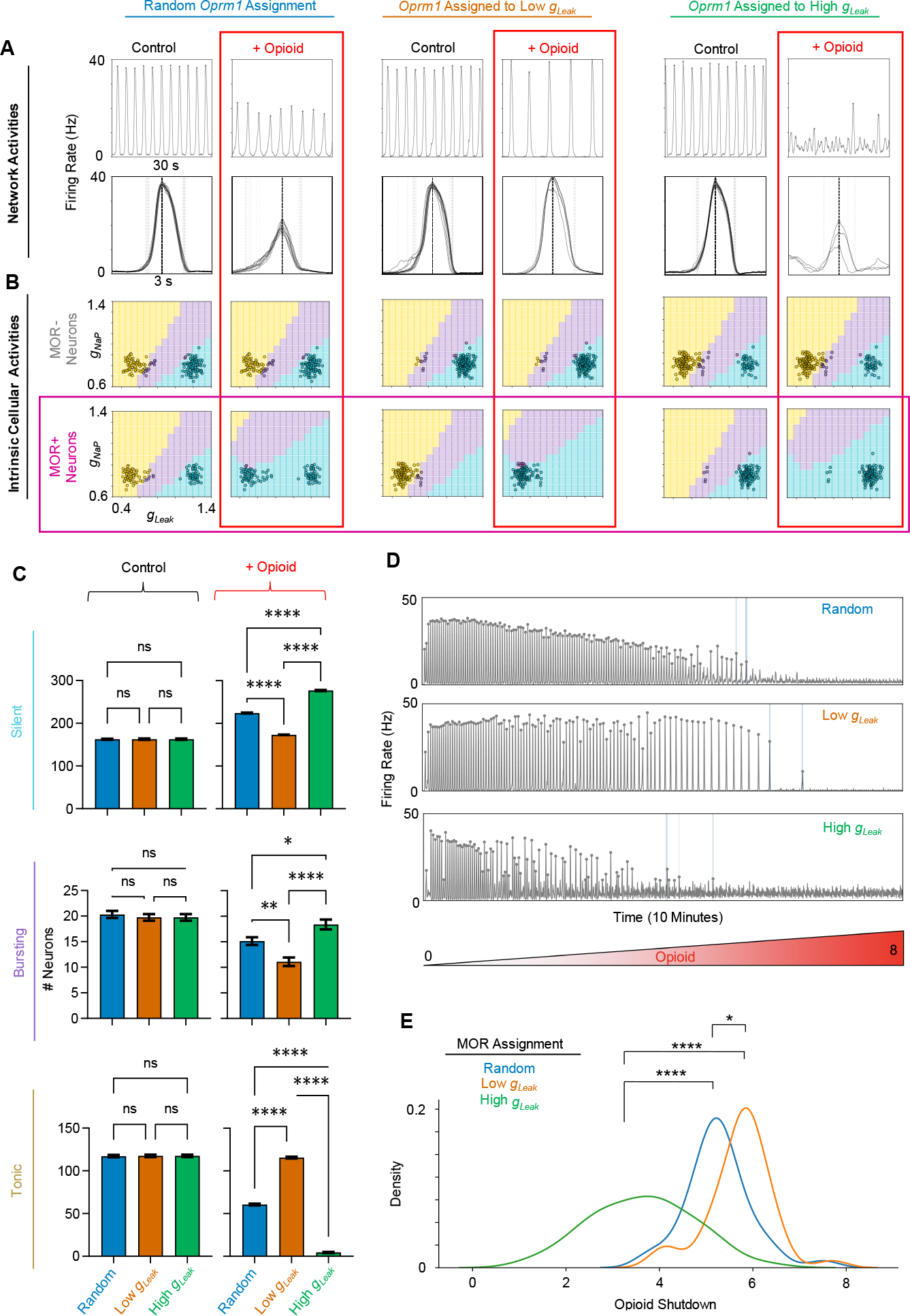
Identity of MOR+ neurons regulates opioid sensitivity. (A) Example rhythms (top) and burst waveforms (bottom) in response to opioids when MOR is assigned randomly (left) or specifically to low *g*_leak_ (middle) or high *g*_leak_ (right) populations. (B) Intrinsic activities of MOR- and MOR+ neurons (open circles) in *g*_NaP_, *g*_leak_ space of the example networks shown in A. C) Quantified number of silent, bursting, and tonic neurons under control conditions and in response to opioid when MOR is assigned randomly or to low/high *g*_leak_ populations (*n* = 40 each, one-way ANOVA with Bonferroni multiple comparisons tests, ns=not significant, *p *<* 0.05, **p *<* 0.01, ****p *<* 0.0001). D) Example network rhythm during opioid ramp (opioid=0-8) with MOR assigned randomly or to low/high *g*_leak_ populations. E) Kernel density estimations showing distributions of opioid shutdown dosages based on the identity of MOR expressing neurons. (*n* = 40, one-way ANOVA with Bonferroni multiple comparisons tests; *p *<* 0.05, ****p *<* 0.0001)

The above results were for a single exemplar network. In Fig. 5C, for each condition, we compared the number of intrinsically silent, bursting, and tonic neurons and how the distributions of these intrinsic activities change in response to opioids across 40 different model networks. As expected (see Fig. 1B), when the identity of MOR+ neurons was randomly assigned, opioids caused many of the low *g*_leak_ MOR+ neurons to transition from tonic/bursting activity to silent, whereas high *g*_leak_ MOR+ neurons were largely unaffected. Under non-random conditions, when all low *g*_leak_ neurons were designated as MOR+, changes in the intrinsic activities within the network were exaggerated such that nearly all intrinsically tonic activity was lost as 92% of the network became intrinsically silent. In contrast, when high *g*_leak_ neurons were designated as MOR+, there were minimal changes in the distribution of intrinsic activities within the networks (Fig. 5C). To further test how the identity of MOR+ neurons may alter how the preBötC network rhythm responds to opioids, we performed simulations ramping up the opioid effect to compare shutdown dosages for networks with MOR identity assigned randomly, or selectively to low or high *g*_leak_ populations. (Fig. 5D, E). Notably, despite a much larger proportion of the network becoming intrinsically silent, networks with low *g*_leak_ neurons designated as MOR+ were more resistant to opioids, than when MOR identity was randomly assigned. On the other hand, the average shutdown dosage was lower when high *g*_leak_ neurons were designated as MOR+, indicating that, despite the minimal effects on the intrinsic activities of the neurons, the network rhythm was substantially more sensitive to opioids under these conditions. These findings support the conclusion that changes in intrinsic cellular activities within the network are not predictive of its sensitivity to opioids (see Fig. 2), but that the distribution of MOR+ expression among preBötC neurons may be an important determinant of how the network responds to opioids.

### 3.5 Modulation of *g*_NaP_ or *g*_leak_ can render the preBötC resistant to opioids

Considering these results, we tested whether manipulations of the intrinsic properties of preBötC neurons may represent a viable strategy to protect the preBötC rhythm from the effects of opioids. Specifically, we tested whether increasing *g*_NaP_ would allow for sustained rhythmogenesis in the presence of relatively high opioid doses as previously hypothesized based on pharmacological experiments *in vitro* (Burgraff et al., 2021). We also tested whether decreasing the leak conductance *g*_leak_ would have a similar protective effect on rhythmogenesis. Rhythmic activity of a representative network under control conditions, in opioid, and during a subsequent 10%, 30%, and 50% increase in *g*_NaP_ are shown in Fig. 6A. Increasing *g*_NaP_ by 30% in the model networks reversed the effects of opioids on burst frequency and amplitude (Fig. 6B). However, recovery of the rhythm by *g*_NaP_ modulation did not restore intrinsic cellular activities to near control. Instead, it was associated with a change in the intrinsic activity of both MOR+ and MOR-neurons from silent to bursting, with little effect on the number of tonic neurons (Fig. 6C, D). Under control conditions, the network was composed of mostly intrinsically tonic and silent neurons (52.3% and 40.7%, respectively). In response to opioid, the proportion of silent neurons increased to 73.7% as MOR+ neurons transitioned from tonic to silent. As *g*_NaP_ was increased, the MOR+ neurons that were originally tonic under control conditions transitioned to bursting. Specifically, when *g*_NaP_ was increased by 30%, 55% of the population entered a *g*_NaP_, *g*_leak_ parameter space that supports intrinsic bursting. Thus, despite recovery of a rhythm with similar frequency and amplitude characteristics following *g*_NaP_ modulation, the number of intrinsically tonic neurons remained reduced, whereas the number of bursting neurons was increased relative to control conditions.

**Figure 6:**
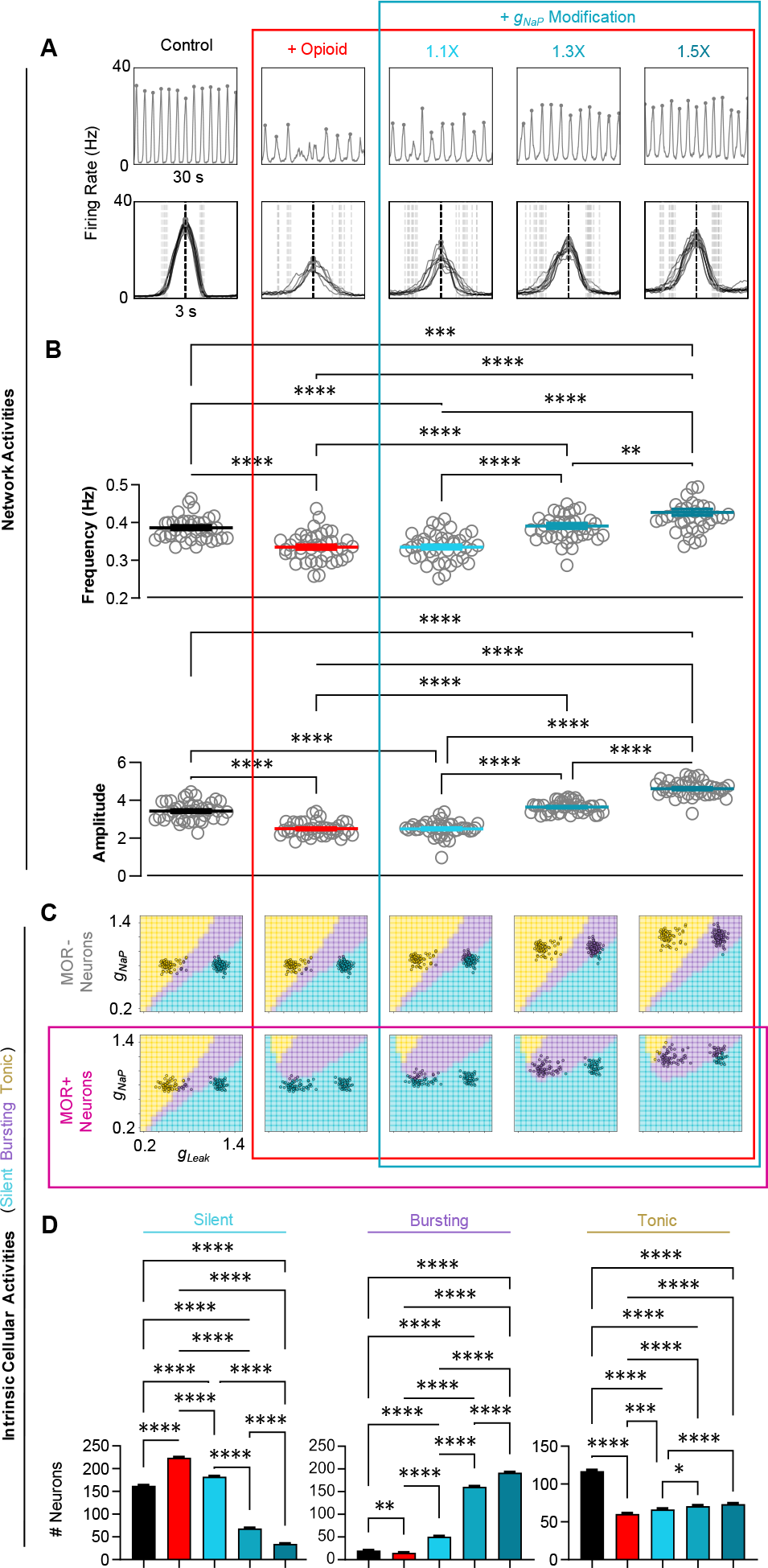
Modulation of *g*_NaP_ renders the network resistant to opioids. (A) Example rhythm and burst waveforms from a network (MOR randomly assigned) in response to opioid and during concurrent modulation of *g*_NaP_ to 110%, 130%, and 150% of control values. (B) Quantified effects on frequency (top) and burst amplitude (bottom) during opioid and *g*_NaP_ modulation (*n* = 40, one-way RM ANOVA with Bonferroni multiple comparisons tests, **p *<* 0.01, ***p *<* 0.001, ****p *<* 0.0001). (C) Changes in the intrinsic activities in *g*_NaP_, *g*_leak_ space of MOR- and MOR+ neurons from the example network shown in A. (D) Quantified changes in the number of silent, bursting, and tonic neurons in response to opioid and subsequent modulation of *g*_NaP_ (*n* = 40, one-way RM ANOVA with Bonferroni multiple comparisons tests, *p *<* 0.05, **p *<* 0.01, ***p *<* 0.001, ****p *<* 0.0001).

We next performed similar simulations during manipulation of *g*_leak_ (Fig. 7). The rhythmic activity of a representative network under control conditions, in opioid, and following a subsequent 10, 30, and 50% reduction in *g*_leak_ is shown in Fig. 7A. In this case, burst amplitude but not frequency could be significantly recovered towards control values (Fig. 7B). This was associated with changes in the intrinsic activities of primarily MOR-neurons (Fig. 7C). When *g*_leak_ was reduced to 70% of control, there was a large increase in the number of bursting neurons, and upon further reduction of *g*_leak_ to 50% of control, these neurons became tonic, leaving only 2.7% of the population as bursting (Fig. 7D). Thus, our model predicts that manipulations that directly or indirectly affect persistent sodium and/or potassium leak conductances may be effective for increasing the resistance of preBötC function to opioids.

**Figure 7:**
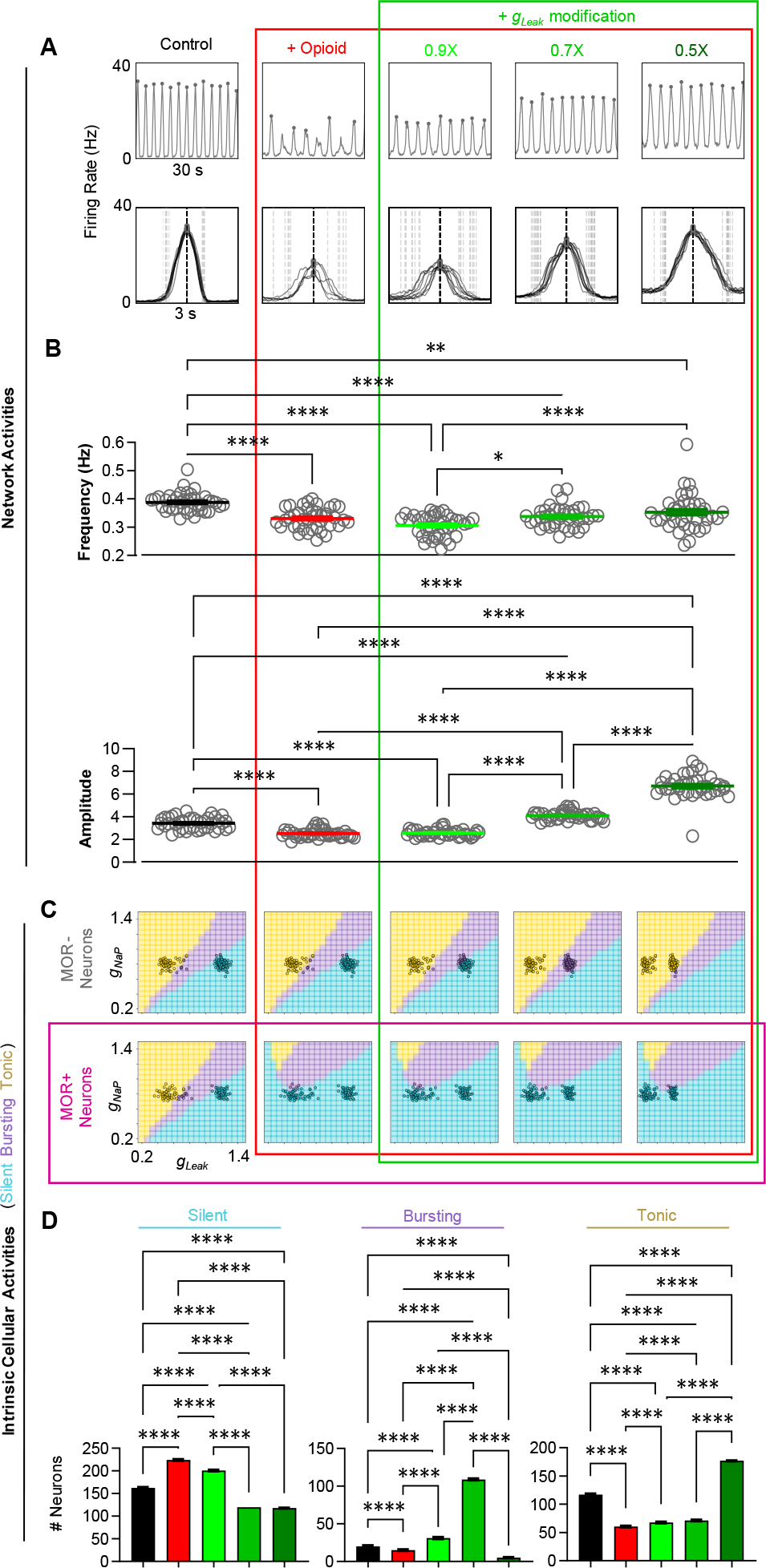
Modulation of *g*_leak_ renders the network resistant to opioids. (A) Example rhythm and burst waveforms from a network in response to opioid and during concurrent modulation of *g*_leak_ to 90%, 70%, and 50% of control values. (B) Quantified effects on frequency (top) and burst amplitude (bottom) during opioid and *g*_leak_ modulation (*n* = 40, one-way RM ANOVA with Bonferroni multiple comparisons tests, **p *<* 0.01, ****p *<* 0.0001). (C) Changes in the intrinsic activities in *g*_NaP_, *g*_leak_ space of MOR- and MOR+ neurons from the example network shown in A. (D) Quantified changes in the number of silent, bursting, and tonic neurons in response to opioid and subsequent modulation of *g*_leak_ (*n* = 40, one-way RM ANOVA with Bonferroni multiple comparisons tests, ****p *<* 0.0001).

## 4 Discussion

The effect of opioids on respiratory function is variable in brain slices *in vitro*, animal models *in vivo*, and in individual humans (Burgraff et al., 2021; Cherny et al., 2001; Dahan et al., 2005; Dahan et al., 2013). Here we adopt a computational model of the respiratory rhythm generator to dissect plausible network topology and cellular properties that contribute to variable respiratory responses to opioids. We leverage computational models that allow us to instantiate networks of the preBötC with connectivity patterns and conductances drawn from random distributions. These networks are statistically indistinguishable on the “macro”-scale; they have the same overall numbers of excitatory and inhibitory neurons, the same numbers of MOR+ and MOR-neurons, the same probability of connections per neuron, and conductance values are drawn from the same distributions. Yet, due to the random assignment of some of these properties, each network differs on the level of individual neurons (nodes), which vary in their exact connectivity patterns and conductance strengths. Surprisingly, this “micro”-level randomness is sufficient to create quite variable responses at the network level to the same stimulus - in this case simulated opioids. We suspect that these differences may contribute to the observed variable responses to opioids seen in experimental preparations (Burgraff et al., 2021). Further, this micro-level variability could, for example, explain how individuals may respond differently to network perturbations despite the preBötC network developing with the same general set of instructions (e.g. genome, transcriptome, axonal targeting mechanisms, etc). While OIRD arises from the effects of opioids on multiple central and peripheral sites (Ramirez et al., 2021), our simulations illustrate how variation in the architecture of the inspiratory rhythm generator could be an important factor underlying the unpredictability of opioid overdose.

The computational approach here allows for directed manipulations that are experimentally intractable. For instance, we are able to ask if the response of the preBötC to opioids depends on MOR being expressed in populations with particular conductance profiles. More concretely, we target the opioid effect directly to neurons that have a particular leak conductance. This leak conductance (*g*_leak_) is an important determinant of whether a neuron is intrinsically “tonic”, “bursting”, or “silent” (Butera et al., 1999b; Del Negro et al., 2002; Koizumi & Smith, 2008; Yamanishi et al., 2018). Surprisingly, introducing MOR selectively to low *g*_leak_ (intrinsically excited neurons with tonic/bursting activity), decreased the response of the network to opioids making the rhythm more resilient. Conversely, introducing MOR selectively to the less excitable population (the high *g*_leak_, quiescent cells), increased the susceptibility of the network rhythm to opioids. We speculate that a robust preBötC rhythm relies on the existence of a population of “recruitable” neurons that are not strongly intrinsically active, but are capable of becoming active with a small amount of synaptic input. When opioids affect neurons in the low *g*_leak_ population, their intrinsic activity is reduced but they remain in the recruitable pool and therefore can continue to participate in the network, allowing the rhythm to continue at higher opioid doses. Conversely, we expect that when opioids further suppress neurons that already have low intrinsic excitability (high *g*_leak_) they are removed from the recruitable pool and unable to participate in network bursts, making coordinated network activity more vulnerable to opioids. When the effect of opioids is randomly targeted to 50% of neurons, the proportion that remains recruitable in the presence of opioid depends on how MOR expression is randomly assigned within the high and low *g*_leak_ populations, contributing to variable opioid responses at the network level.

Network connectivity is difficult to study and manipulate experimentally. Thus, computational models, where the number and strength of all connections between every neuron are known, can be an important tool to provide “proof of concept” insights into how network topology can influence network function and determine its response to perturbations. We took advantage of this by performing correlation analysis to better understand how the number of connections between certain subgroups of preBötC neurons may predict how susceptible the network is to opioids. These analyses revealed that, in general, when neurons that do not respond to opioid (MOR-) are more interconnected and receive less inhibitory input, the network is more resistant to opioids. We suspect that this connectivity configuration may allow the network of MOR-neurons to remain rhythmogenic even when very few opioid sensitive (MOR+) neurons are able to contribute to network function. In another analysis, we scaled the number of connections within the network without altering total synaptic strength, which consistently increased the robustness of the network to opioids. Because opioids weaken the pre-synaptic strength of excitatory interactions (Baertsch et al., 2021), we anticipate that networks with lower numbers of connections become “fractured” into isolated sub-networks when opioid-induced weakening of synapses impairs the network’s ability to effectively recruit portions of the population. Indeed, the preBötC rhythm *in vitro* has a higher proportion of failed bursts with low amplitude in response to opioids (Baertsch et al., 2021; Phillips & Rubin, 2022). In networks with more connections, activity more consistently propagates to all neurons (Kam et al., 2013), efficiently recruiting the whole population despite the effect of opioids on synaptic transmission. This could also contribute to the variable opioid responses observed in *in vitro* experiments since both within and across labs where the creation of rhythmic brain stem slices invariably samples slightly different portions of the preBötC population that may be more or less densely connected (Baertsch et al., 2019; Ruangkittisakul et al., 2014). Although these simulations illustrate that network topology could be an important determinant of opioid sensitivity, because connection density and patterns are considered “fixed” properties of the network, at least on short time scales, manipulation of network topology is an unlikely avenue for therapeutic interventions. In contrast, the strength of existing excitatory synaptic connections can be pharmacologically altered acutely via e.g. ampakines, which may render the preBötC less vulnerable to opioids and shows promise as an intervention for OIRD (Ren et al., 2006; Sunshine & Fuller, 2021; Xiao et al., 2020).

The intrinsic activity of preBötC neurons is determined by multiple interacting cellular properties (Ramirez et al., 2012). Not all are known and not all can be incorporated into our simplified model network. Yet, like many other computational studies (Lindsey et al., 2012), the interaction between *g*_leak_ and *g*_NaP_ determines intrinsic activity in our model and is sufficient to capture the silent, bursting, or tonic phenotypes of preBötC neurons. Both *g*_leak_ and *g*_NaP_ contribute to cellular excitability (resting membrane potential), and the voltage-dependent properties of *g*_NaP_ allow some neurons with appropriate *g*_leak_ to exhibit intrinsic bursting or “pacemaker” activity (Koizumi & Smith, 2008). Whether such neurons with intrinsic bursting capabilities have a specialized role in network rhythmogenesis is a matter of ongoing debate (da Silva et al., 2023; Feldman & Del Negro, 2006; Ramirez & Baertsch, 2018a; Ramirez & Baertsch, 2018b; Smith et al., 2000) that we do not address here. Instead, we aimed to understand how opioids alter the intrinsic activities of preBötC neurons. In the model network, opioids reduce the number of neurons with intrinsic bursting or tonic activity and increase the number of silent neurons. To our surprise, the extent of these changes was not a significant predictor of the network response to opioids. This suggests that the intrinsic activity of a given neuron may not be representative of its contribution to network function, and that other factors, such as those discussed above, play more substantial roles in determining how the preBötC responds to opioids. Although network differences due to random sampling of *g*_leak_ and *g*_NaP_ from set distributions were not a significant factor driving variable opioid responses, we found that scaling the distribution of *g*_NaP_ or *g*_leak_ across the whole population did alter the sensitivity of model networks to opioids. Interestingly, manipulation of *g*_NaP_ was more effective since a 30% increase in *g*_NaP_ was sufficient to restore both frequency and amplitude of the rhythm, whereas effects were more specific to burst amplitude following a 30% decrease in *g*_leak_. Unlike network topology, intrinsic conductances that regulate cellular excitability and activity are not “fixed” but are dynamic and can be modified by conditional changes in e.g. neuromodulators and ion concentrations (Ramirez et al., 2012; Rybak et al., 2007) and are also more amenable to pharmacological manipulations (Bedoya et al., 2019; Burgraff et al., 2021; Verneuil et al., 2020). Thus, further experimental investigation of these approaches is warranted as they may hold promise as potential therapeutic strategies to protect against opioid-induced failure of preBötC network function.

## Notes

https://github.com/glomerulus-lab/prebot-opioid-model

### Competing Interest Statement

The authors have declared no competing interest.

### Summary of Updates

Revisions to Figures, Intro, and Conclusions

